# Microdissection and culturing of adult lateral entorhinal cortex layer II neurons from APP/PS1 Alzheimer model mice

**DOI:** 10.1101/2022.12.05.519179

**Authors:** Katrine Sjaastad Hanssen, Menno P. Witter, Axel Sandvig, Ioanna Sandvig, Asgeir Kobro-Flatmoen

## Abstract

**Background:** Primary neuronal cultures enable cell-biological studies of Alzheimer’s disease (AD), albeit typically non-neuron-specific. The first cortical neurons affected in AD reside in layer II of the lateralmost part of the entorhinal cortex, and they undergo early accumulation of intracellular amyloid-β, form subsequent tau pathology, and start degenerating pre-symptomatically. These vulnerable entorhinal neurons uniquely express the glycoprotein reelin and provide selective inputs to the hippocampal memory system. Gaining a more direct access to study these neurons is therefore highly relevant.

**New method:** We demonstrate a methodological approach for microdissection and long-term culturing of adult lateral entorhinal layer II-neurons from AD-model mice.

**Results:** We maintain adult microdissected lateralmost entorhinal layer II-neurons beyond two months in culture. We show that they express neuronal markers, and that they are electrophysiologically active by 15 days *in vitro* and continuing beyond 2 months.

**Comparison with existing methods:** Primary neurons are typically harvested from embryonic or early postnatal brains because such neurons are easier to culture compared to adult neurons. Methods to culture adult primary neurons have been reported, however, to our knowledge, culturing of adult entorhinal subregion-specific primary neurons from AD-model animals has not been reported.

**Conclusions:** Our methodological approach offers a window to study initial pathological changes in the AD disease-cascade. This includes the study of proteinopathy, single-neuron changes, and network-level dysfunction.

**Highlights:**

- We microdissect and culture neurons from layer II of the lateralmost part of the entorhinal cortex from adult AD model mice and littermate controls
- These entorhinal neurons self-organize into networks, express reelin, NeuN and intracellular amyloid-β.
- The neurons are electrophysiologically active by day 15 in culture and remain viable beyond two months.

## 1. Introduction

For more than two decades, cultured cells have been widely used to model aspects of the neurodegenerative cascade inherent to Alzheimer’s disease (AD) (Trinchese et al., 2004), and constitute an indispensable tool to complement *in vivo* models (Belle et al., 2018). Cellular culture models typically make use of established cell lines of rodent or human origin, or they rely on harvesting primary neurons. The latter are typically obtained from embryonic or early postnatal cortical tissue from rats or mice (Calvo-Rodríguez et al., 2017; Kaech et al., 2006; Konings et al., 2021; Roos et al., 2021; Sahu et al., 2019). However, AD affects subsets of mature neurons, and we may therefore gain insights also by studying cultures of fully developed neurons, for example by harvesting these from *adult* AD-model animals. This approach not only avoids possible confounds from developmental factors operating in embryonic and early postnatal neurons, but may also offer a better representation of how neurons respond to AD-relevant pathologies in the actual disease. Methods to extract and culture adult primary neurons from mice (Brewer et al., 2007; Eide et al., 2005; Varghese et al., 2009) and rats (Brewer, 1997; Brewer et al., 2007) have previously been reported, however, work on cultures of subregion-specific neurons from adult AD-model animals is lacking.

Immunohistochemical studies (Braak et al., 1991; Gomez-Isla et al., 1996; Kordower et al., 2001) as well as sophisticated live human imaging methods (Holbrook et al., 2020; Kulason et al., 2019; Olsen et al., 2017; Tran et al., 2022; Tward et al., 2017) show that cortical AD-related pathological changes predominantly start in layer II (LII) neurons located in the lateralmost part of the entorhinal cortex (EC), i.e., close to the collateral sulcus. In rodents, equivalent neurons are situated in the cytoarchitectonically defined lateral entorhinal cortex (LEC), and, even though exact borders for a corresponding cytoarchitectonic subdivision of human EC have not yet been decisively established, the same LII neuron-types are present in the lateralmost EC of rodents as those in the lateralmost EC of humans (reviewed in (Kobro-Flatmoen et al., 2019)). Such ECLII-neurons exhibit early AD-related alterations in firing properties (Marcantoni et al., 2014), tend to form tangle pathology prior to other cortical areas (Berron et al., 2021; Braak et al., 1991), and start to die already at preclinical stages of AD (Gomez-Isla et al., 1996; Kordower et al., 2001).

In humans as well as rodents, AD vulnerable ECLII-neurons are unique in expressing reelin (Re+) (Kobro-Flatmoen et al., 2019), which is a large glycoprotein that promotes synaptic plasticity and, of particular relevance to AD, inhibits glycogen synthase kinase 3β-mediated phosphorylation of tau. Work on rodents has established that levels of reelin in such Re+ ECLII-neurons align along a gradient where the lateralmost neurons express high levels of reelin, while those located increasingly further medially express successively lower levels of reelin (Kobro-Flatmoen et al., 2016). Intriguingly, such Re+ ECLII-neurons are highly prone to accumulating intracellular amyloid-β42 (Aβ42), and this accumulation of Aβ42 takes the form of a pattern which aligns along the same gradient as that for reelin (Kobro-Flatmoen et al., 2016). Recent experimental work established that in Re+ ECLII-neurons, Aβ42 directly interacts with reelin. Furthermore, levels of Aβ42 are dependent on levels of reelin, and selectively lowering levels of reelin in these neurons results in a concomitant lowering of Aβ42-levels (Kobro-Flatmoen et al., 2022).

Using adult APP/PS1 mice (Radde et al., 2006), we here present a methodological approach to microdissect and culture neurons from the lateralmost part of LEC layer II (lLEC-LII). Figure 1 shows an overview of our experimental setup. Our work is based on mice at ages ranging from postnatal day (P) 48 to P60, thus representing the brain with a fully developed EC (Donato et al., 2017; Semple et al., 2013). We find that cultured adult lLEC-LII-neurons form connections within the two first weeks and remain viable beyond two months. We confirm the lLEC-LII-specific nature of our cultured neurons by showing that they co-express NeuN and reelin. We also find that reelin expression colocalizes with that of Aβ, indicating that the tendency of such neurons to express Aβ (Kobro-Flatmoen et al., 2016) is retained when they are cultured. The lLEC-LII-networks show spontaneous neuronal activity by two weeks post-seeding and remain active beyond two months. Our approach may thus represent a valuable tool for studying very early proteinopathy, and offers neuron- and network specific modelling of AD pathology *in vitro*.

**Figure 1.**
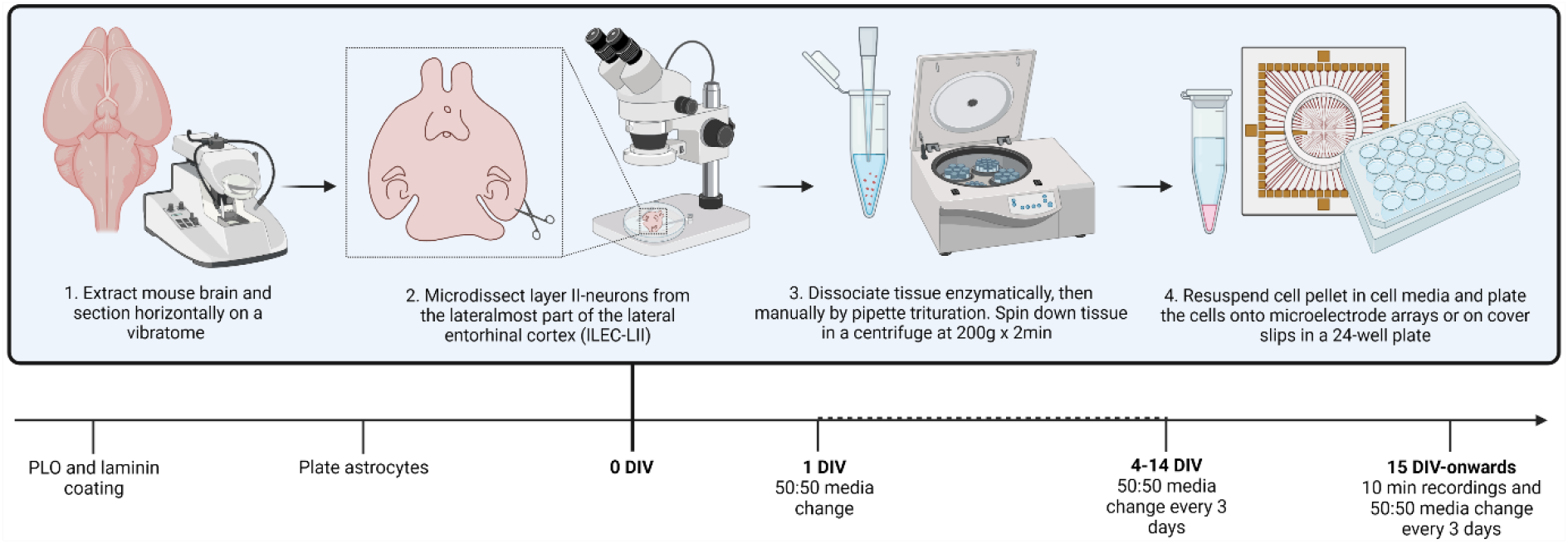
Overview of experimental setup. Before plating of the lLEC-LII-neurons, we coated the culture wells with Poly-L-Ornithine (PLO) and laminin. The following day we added a monolayer of rat cortical astrocytes. On day 0, we extracted and horizontally sectioned APP/PS1 mouse brains, and then microdissected strips of lLEC-LII-neurons under a stereoscope. The strips were dissociated enzymatically and then manually triturated with a 100 µL pipette, and then spun down in a centrifuge at 200g x 2min. The cell pellet was resuspended in 100 µL cell media before plating. At 1 DIV (days *in vitro*) half the media was changed; this was subsequently repeated every 3 days. From 15 DIV we started recording spontaneous activity. DIV; days *in vitro*. lLEC-LII; lateralmost lateral entorhinal cortex layer II.

## 2. Materials and methods

### 2.1. Animals

To develop the cell culture method, we used a total of 48 adult mice, including APP/PS1 model mice and littermate controls, ranging in age from postnatal day 28 (P28) to P115, including 25 males and 23 females. Of these, we collected data from 9 animals (7 males, 2 females; Supplementary Table A.1). All our work complies with the ARRIVE guidelines (Percie du Sert et al., 2020) and the protocol was approved by The Norwegian Food Safety Authority (FOTS ID 8433 and ID 22321), according to the <u>EU Directive 2010/63/EU</u>for animal experiments. Animals were bred at the Comparative Medicine Core Facility (CoMed) at the Norwegian University of Science and Technology (NTNU) and were held in standard lab cages (up to 5 animals per cage), in temperatures of 22 ± 2°C, and kept at a light/dark cycle of 12:12 hours, with access to food and water *ad libitum*. Minimum one week before experiments, animals were transferred to the animal facility at the Kavli Institute for Systems Neuroscience, Centre for Neural Computation, Norwegian University of Science and Technology (NTNU), where brain extractions and microdissections were performed. Subsequent processing and experiments were carried out at the Department of Neuromedicine and Movement Science, NTNU.

#### 2.1.1. APP/PS1 mouse model and genotyping

The APP/PS1 mouse model is a transgenic familial AD-model, with a C57BL/6J genetic background, co-expressing mutated human amyloid precursor protein (KM670/671NL) and mutated human presenilin 1 (L166). In the model, both mutated proteins are driven by expression cassettes under the Thy1 promoter, which causes accumulation of Aβ with extracellular cortical deposits appearing from around 6 weeks of age (Radde et al., 2006). The animals used in our experiments were genotyped for expression of the transgene by quantitative PCR (qPCR) using the KAPA Mouse genotyping kit (Merck, Cat# MGKITKB-KK7301), with primer sequence (5’-3’) APP-F (GATTCCGACATGACTCAGG) and APP-R (CTTCTGCTGCATCTTGGACA).

### 2.2. Preparation of culture plates, coating, and seeding of astrocytic monolayer

For electrophysiological recordings we used microelectrode arrays (MEAs; 60EcoMEA-Glass-gr, Multichannel Systems, MCS, Reutlingen, Germany), while for immunocytochemistry (ICC) we used round coverslips (13mm) in a 24-well plate. Additionally, ICC was done on one MEA. Cleaning, sterilization, and impedance measurement of the MEAs was done as recommended by the manufacturer (<u>Microelectrode Array (MEA) Manual</u>). The culture surfaces (MEAs and glass coverslips) were first coated with 400 μL Poly-L-Ornithine (PLO) in an incubator at 37°C with 5% CO_2_ for 2 hours. Subsequently, all PLO was discarded, and the surface was rinsed gently three times with sterile Milli-Q-water before we added 400 μL laminin solution. We then incubated the culture vessels at 37°C with 5% CO_2_ for 2 hours. The laminin solution was then discarded, and 200 μL fresh pre-warmed astrocyte media was incubated in the culture plates at 37°C with 5% CO_2_ for ∼10 min, before plating ∼1.500 Rat Primary Cortical Astrocytes per culture well (solutions are listed in Table 1). The astrocytes were plated 24-48 hours prior to plating the lLEC-LII neurons. Through extensive pilot experimentation, we concluded that a monolayer of astrocytes is crucial for the attachment and viability of adult lLEC-LII neurons, and a complete lack of neuronal attachment resulted when astrocytes were plated either simultaneously or following that of the neurons.

**Table 1.** Composition of solutions.

| Dissection medium (HABG)* | Cell medium (growth medium) | Laminin solution |
| --- | --- | --- |
| • Hibernate™-A Medium<br>• 2% B27™ Plus Supplement<br>• 0.5 mM GlutaMAX™ Supplement (2mL/L)<br>• 1% Penicillin-Streptomycin | • Neurobasal Plus Medium<br>• 2% B27™ Plus Supplement<br>• 0.5 mM GlutaMAX™ Supplement (2mL/L)<br>• 10% Fetal Bovine Serum<br>• 1% Penicillin-Streptomycin<br>• 1:1000 RHO/ROCK pathway inhibitor<br>• 1:1000 BDNF Human Recombinant | • L15 medium<br>• 2.5 % Sodium Bicarbonate<br>• 1,66 % Mouse laminin |
|  |  | Astrocyte medium** |
|  |  | • 85% DMEM (high glucose)<br>• 15% Fetal Bovine Serum<br>• 1% Penicillin-Streptomycin |

\*Same media used for microdissection and centrifugation of neurons. \*\*Media recommended by the manufacturer of rat primary cortical astrocytes (Gibco™, #N7745-100).

### 2.3. Dissection of brain tissue

Table 2 lists all media components used for the microdissection and culture. All tools were sterilized by autoclaving before use. Mice were deeply anesthetized using isoflurane gas (Abbott Lab, Cat# 05260-05) and then decapitated using surgical scissors (FST, Cat# 14007-14). Subsequently, the head was quickly immersed in 70% ethanol followed by sterile phosphate buffered saline, before being submersed in ice cold Hanks Balanced Salt Solution (HBSS) in a sterilized glass dish resting on ice. While fully submerged, we cut and peeled away the scalp, and then used fine scissors (FST, Cat# 14568-09) to open the skull along the midline, before prying each resulting half of the skull aside using the same scissors along with a pair of S…T forceps (FST, Cat# 00108-11), thus exposing the brain. A spatula (RSG Solingen, Cat# 231-2262) served to sever the cranial nerves, allowing us to gently lift out the brain and transfer it to a sterile cup containing ice cold HBSS. Immediately before mounting the brain, we took it out of the HBSS in order to slice off the cerebellum plus ∼300 μm of the brains’ dorsal surface, which gave us a flat surface for better attachment. Subsequently, that (dorsal) surface of the brain was glued to the specimen disc using Loctite 401 superglue (Henkel-Adhesives). Abutting the caudal end of the brain we glued on a supportive barrier of freshly made AGAR (VWR; Cat# 20767.298) for structural support vis a vis the direction traversed by the blade. The specimen disc containing the brain and the supportive AGAR was then submerged into the vibratome container containing ice cold HBSS. We sectioned in the rostral-to-caudal direction at a thickness of 300 μm, setting the blade at a frequency of 5.3 Hz with a speed of 8mm/s. By using sterilized plastic pipettes cut to shape, each section was transferred to separate wells in a sterile 24-well plate resting on ice, containing a solution of Hibernate-A medium, 2% B27 Plus Supplement, and 2.5mM/mL GlutaMAX Supplement (HABG; (Brewer et al., 2007)), in addition to 1% penicillin streptomycin. A sterile glass petri dish, placed up-side-down on top of a pool of ice at the stage of a stereoscope (SteREO Discovery V8; Zeiss), served as our dissection platform. One-by-one we then transferred each section from the 24-well plate onto the surface of the petri dish, which, viewed under the stereoscope using cross polarized optic fibre lights, allows for sufficient contrast to recognize anatomical landmarks. We then microdissected lLEC-LII as detailed in the subsequent section.

**Table 2.** Reagents, equipment, materials, and antibodies.

| Reagents |  |
| --- | --- |
| Accutase Cell Dissociation Medium | (Thermo Fischer Scientific, Cat# A1110501) |
| AGAR | (VWR; Cat# 20767.298) |
| Anti-Fade Fluorescence Mounting Medium – Aqueous, | (Abcam, Cat# ab104135) |
| B-27™ Plus Supplement | (Gibco™, Cat# A3582801) |
| Brain derived neurotrophic factor [BDNF Human Recombinant] | (Neurotrophins, Cat# 450-02) |
| Dulbecco's Phosphate Buffered Saline | (Thermo Fischer Scientific, Cat# 14040-117) |
| DMEM (high glucose) | (Gibco™, Cat# 11995065) |
| Fetal Bovine Serum (FBS) | (Sigma Aldrich, Cat# 12106C-100ML) |
| GlutaMAX™ Supplement | (Gibco™, Cat# 35050061) |
| Goat Serum | (Sigma-Aldrich, Cat# G9023) |
| Hanks Balanced Salt Solution (HBSS) | (Thermo Fischer Scientific, Cat# 88284) |
| Hibernate™-A Medium | (Thermo Fischer Scientific, Cat# A1247501) |
| Hoechst 33342 Trihydrochloride | (Sigma-Aldrich, Cat# 14533) |
| Isoflurane | (Abbott Lab, Cat# 05260-05) |
| Laminin Mouse Protein, Natural | (Thermo Fischer Scientific, Cat# 23017015) |
| L15 Medium | (Sigma-Aldrich, Cat# L5520) |
| Neurobasal Plus Medium | (Gibco™, Cat# A3582901) |
| Neuronal isolation enzyme Papain | (Thermo Fischer Scientific, Cat# 88285) |
| Paraformaldehyde | (Sigma-Aldrich, Cat# P6148) |
| Penicillin Streptomycin (PS) | (Gibco™, Cat# 15070063) |
| Poly-L-Ornithine (PLO) | (Sigma-Aldrich, Cat# P3655-50mg) |
| Rat Primary Cortical Astrocytes | (Gibco™, Cat# N7745-100) |
| Sodium Bicarbonate | (Sigma Aldrich, Cat# S5761) |
| StemPro™ Accutase™ Cell Dissociation Enzyme | (Gibco™, Cat #A1110501) |
| Trypan Blue Stain | (Sigma-Aldrich, Cat# T10282) |
| Y-27632 (Dihydrochloride) RHO/ROCK pathway inhibitor (RI) | (STEMCELL™ Technologies, Cat# 72302) |
| Equipment and materials |  |
| BrandTech Accu-jet pro Pipette Controller | (BrandTech Scientific, Cat# 26330) |
| Corning 24 well plates | (VWR, Cat #10062-896) |
| Countess II Automated cell counter | (Thermo Fischer, Cat #AMQAX1000) |
| EVOS M5000 Imaging System | (Thermo Fischer Scientific, Cat #AMF5000) |
| Falcon tube (15 mL) | (Merck, Cat #CLS430791) |
| Surgical Scissors | (FST, Cat# 14007-14) |
| Fine Scissors | (FST, Cat# 14568-09) |
| RSG Solingen Spatulae | (VWR, Cat# 231-2262) |
| S&T Forceps | (FST, Cat# 00108-11) |
| Stereoscope | (SteREO Discovery V8; Zeiss) |
| Vannas Spring Scissors | (FST, Cat# 15018-10) |
| Vibrating blade microtome | (Leica Biosystems, Cat# 14047235613) |
| **Primary and secondary antibodies** |  |
| Anti-Reelin Antibody (1:1000) | (Abcam, Cat #ab78540) |
| Anti-NeuN Antibody (1:1000) | (Sigma-Aldrich, Cat# ABN90P) |
| Oligomer A11 Polyclonal Antibody (1:1000) | (Invitrogen, Cat# AHB0052) |
| Alexa Fluor™ 488 Goat-anti-Mouse IgG | (Thermo Fischer Scientific, Cat #A-11001) |
| Alexa Fluor™ 546 Goat-anti-Guinea Pig IgG | (Thermo Fischer Scientific, Cat #A-11074) |
| Alexa Fluor™ 647 Goat-anti-Rabbit IgG | (Thermo Fischer Scientific, Cat #A32733) |

#### 2.3.1. Microdissecting entorhinal cortex layer II-neurons from the lateral domain

EC is a six-layered periallocortex. Rodent EC contains two major cytoarchitectonic subdivisions that are anchored to specific connectivity patterns, known as the medial entorhinal cortex (MEC) and the lateral entorhinal cortex (LEC) (Insausti et al., 1997). For reasons explained in the introduction, we specifically targeted the *lateralmost part of LEC layer II* (lLEC-LII), i.e., the part close to the rhinal fissure, by taking advantage of a set of characteristics that are recognizable on fresh sections when viewed under a stereoscope and illuminated by cross polarized light. Thus, we separated LEC from MEC by identifying and tracing the characteristic layer V neurons of MEC, which align to give the appearance of radial columns, a feature that abruptly disappears to mark the medial border of LEC. We then deliberately placed our initial cut a distance away from this border to ensure targeting of lLEC-LII. On the other side, literally, as one moves into the most lateral domain of LEC, big fan and pyramidal neurons locate superficially in LII (sometimes referred to as layer IIa; these neurons stain positive for reelin). These LII-neurons are separated from a more deeply situated layer of pyramidal neurons (IIb, calbindin-positive) by a cell-free zone that is only present towards the lateral extreme. This feature, together with the notably smaller superficial neurons residing in the perirhinal- and postrhinal cortices, which form the anterior and dorsal borders of LEC, is sufficient to reliably delineate and microdissect lLEC-LII neurons (Kobro-Flatmoen et al., 2019). We cut strips of lLEC-LII tissue using Vannas Spring scissors (FST, Cat #15018-10), immediately transferred the strips to 15 mL Falcon tubes containing HABG (see above, and Table 1), and immersed these tubes in crushed ice. To safeguard our ability to target the region of interest, we used sections from one mouse and mounted and stained these sections using cresyl violet (Fig. 2). This, along with immunocytochemistry (ICC; see below), confirmed our ability to consistently target and extract lLEC-LII neurons.

**Figure 2.**
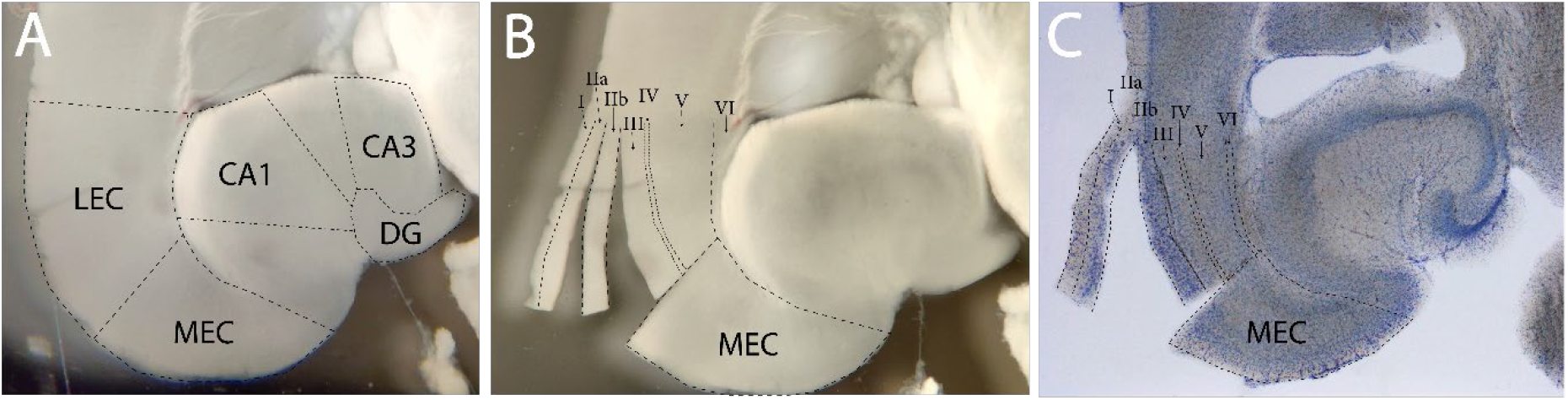
Microdissecting lLEC-LII neurons. **(A)** Horizontal brain section cut at 300 μm. Salient neuroanatomical features are recognizable when viewed in a stereoscope under cross-polarized light, using optic lamps. **(B)** Same brain section as in (A) with tissue delineated (dashed lines) and LEC layers IIa and IIb cut and slightly spread apart. **(C)** Same brain section as in (A-B) stained with Cresyl Violet. Note that layer IIb has now been pushed against LIII, such that only LIIa points out towards the left side. Furthermore, due to drying and coverslipping, the section is shrunk and flattened, and the delineations are adjusted accordingly. Abbreviations: LEC, lateral entorhinal cortex; lLEC, lateralmost lateral entorhinal cortex; MEC, medial entorhinal cortex; DG, dentate gyrus; CA3, cornu ammonis 3; CA1, cornu ammonis 1; LII, layer II.

### 2.4. Processing and culturing of lLEC-LII neurons

To remove potential debris from the microdissection, we discarded HABG from the 15 mL falcon tubes and rinsed the tissue strips by adding and removing 3 mL cold Hibernate-A three times, letting the tube rest in ice for 20 seconds in between each rinse. We then transferred the tissue into a sterile 1.5 mL Eppendorf tube containing 250 μL neuronal isolation enzyme Papain and incubated the tissue in a standard humified incubator (37°C, 5% CO_2_, 20% O_2_) for 15 minutes. Subsequently, we discarded the dissociation enzyme and added 100 μL pre-warmed HABG and manually triturated the tissue by running it ten times through a 100 μL pipette at a rate of about 45 seconds. The resulting cell suspension was transferred to a 15 mL Falcon tube containing 2.9 mL pre-warmed (37°C) HABG, to give a total of 3 mL, and then centrifuged at 200x g for 2 minutes at 21°C (room temperature). Subsequently, we discarded all HABG, added 100 μL cell media (see Table 1), and completed the dissociation by again triturating the tissue by running it through a 100 μL pipette ten times, while taking care to ensure no air bubbles reached the pipette. A cell count was performed using a Countess™ Automated cell counter (Thermo Fischer, Cat #AMQAX1000). For each separate culture (MEAs or 24-well plates) we used neurons from two animals, which gave between 140,000 and 170,000 cells/well. The neurons (cell suspension) were plated in 100 μL cell media and incubated for 2 hours, then 800 μL of fresh pre-warmed (37°C) cell media was added to give a total of 1 mL cell media. The cultures were incubated for 24 hours at 37°C (5% CO_2_, 20% O_2_). Then, 50% of the cell media was replaced with fresh pre-warmed (37°C) cell media. In cases of excess debris in the cultures 24 hours after plating, we carried out extra rinses by aspirating 90% of the media, and then tilted the cultures slightly while slowly adding 500 μL cell media such that the culture was gently flushed, before we added new media.

We supplemented the cultures every third day by aspirating half the cell media and replenishing it with fresh, pre-warmed media (37°C); this frequency of media changes was determined to be optimal by extensive experimentation (Supplementary fig. A.1). Degenerating and dying neurons were readily identifiable by observing their retracting neurites and detaching cell bodies. On the days of electrophysiological recordings, media changes were made after the recording was completed. Imaging and media changes were done within a time window of <3 minutes, as longer time periods outside the incubator will affect the cultures’ pH levels and induce cytotoxicity.

### 2.5. Immunocytochemistry (ICC) and imaging

ICC was carried out as follows. We aspirated all media from the cultures before adding 500 μL Dulbecco’s Phosphate Buffered Saline (DPBS) to clear away debris. Then we fixed the cultures by applying 4% freshly depolymerized paraformaldehyde for 15 minutes, followed by 10×3 minute rinses in DPBS. To block unspecific antigens, we incubated the cultures in a solution of DPBS containing 3% goat serum and 0.1% Triton-X, for 1 hour at room temperature. The cultures were then incubated overnight with primary antibodies (see Table 1) in a solution of DPBS containing 1% goat serum, on a shaker at 4°C. Then the cultures were rinsed three times in DPBS, immediately before incubation in DPBS containing secondary antibodies at room temperature for 2 hours (see Table 1). Hoechst dye was added (1:10.000) for the last 10 minutes of the secondary antibody incubation time. Images were acquired with an EVOS M5000 microscope connected to a LED light source and using an Olympus 20x/0.75 NA objective (N1480500), with the following filter sets: DAPI (AMEP4650), CY5 (AMEP4656), GFP (AMEP4651) and TxRed (AMEP4655). For image processing we used Zen 2.6 (blue edition), along with contrast adjustments in Adobe Photoshop 2020.

### 2.6. Electrophysiological recordings

We recorded electrophysiological activity from two cultures on MEAs using a Multichannel MEA2100 recording system (Multichannel Systems, MCS, Reutlingen, Germany). The stage temperature was set to 37°C, which minimizes thermal stress, and we used EcoMEAs covered with a MEA-MEM-Ring (Multichannel Systems, MCS, Reutlingen, Germany) to avoid contaminants and media evaporation. After removal from the incubator, the cultures were left to rest for ∼3 minutes on the stage before spontaneous electrophysiological activity was recorded for 10-minute epochs. This recording time was chosen as our system does not maintain stable CO2-levels, such that time out of the incubator will affect pH levels and thus cell-viability (Lloyd et al., 2015). Data sampling was collected at a rate of 10 kHz/channel. We converted raw data to an .h5 Hierarchical Data Format file using the Multichannel DataManager (V.1.14.4.22018) system and imported it to MatLAB2020a for analyses. Spontaneous neuronal network activity was thus recorded at the following timepoints (DIV): 15, 18, 24, 27, 30, 33, 39, 48, 54, 57, 60, 62 and 69 (n=2; MEA 1 and MEA 2).

### 2.7. Data analysis

For spike detection we used the Precise Timing Spike Detection (PTSD) algorithm developed by (Maccione et al., 2009). To reduce artifacts from the raw data we used a fourth order Butterworth-filter with a high pass cutoff frequency at 300 Hz and a low pass cutoff frequency at 3000 Hz. We additionally used a Notch filter to exclude electrical noise at 50 Hz from the mains hum. A signal was defined and recorded as a spike when reaching beyond a differential threshold of 7.5 times the standard deviation of the background activity. Firing rate and burst propensity (a measure of fraction of spikes in networks bursts) were analysed using an in-house made MATLAB script, reported previously (Fiskum et al., 2021). A network burst was identified when a binned spike distribution of 50ms bins exceeded a threshold of firing rate of 7.5 standard deviations and more than 20% of all active electrodes in the recording fired.

## 3. Results

### 3.1 Microdissection and culturing of lLEC-LII neurons

Our procedure to microdissect and culture adult lLEC-LII neurons is outlined in Figure 1. Each culture consisted of cells from two animals, and, at the time of plating, contained from 140,000 to 170,000 cells as estimated by a Countess II Automated cell counter (Thermo Fischer, Cat #AMQAX1000). Putative neurons attached and began extending neurites already by 2 hours *in vitro* (Fig. 3A). By 14 DIV, we observed a pronounced growth of neurites, and many of these had already formed structural networks (Fig. 3 B-D). Such networks remained viable beyond two months, at which point we terminated the cultures to establish cell-identities by way of ICC (Fig. 3E-N). We found our cultures to be rich in cells that stained double-positive for NeuN and reelin, providing strong support that we in fact successfully acquired Re+ lLEC-LII-neurons (Fig. 3; Supplementary Fig. A.2). We furthermore found that the Re+ neurons co-expressed oligomeric Aβ (Fig. 3), in line with the known strong intracellular expression of Aβ in such neurons (Kobro-Flatmoen et al., 2016).

**Figure 3.**
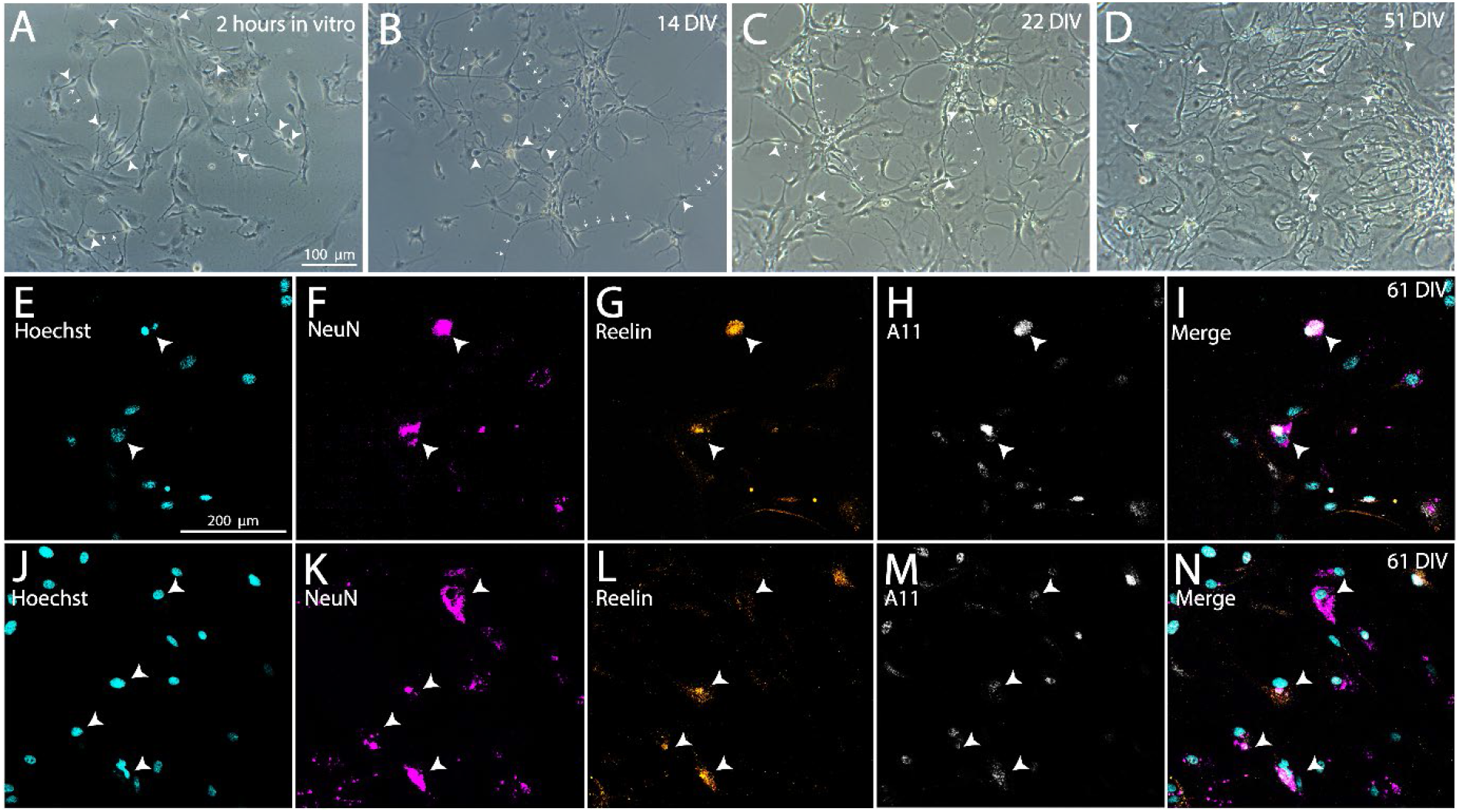
Adult microdissected lLEC-LII-neurons self-organize in culture and express selective markers. **(A-D)** Phase-contrast images of lLEC-LII neurons from the same culture at 2 hours (A), 14 days (B), 22 days (C), 51 days (D) *in vitro*. Arrowheads indicate neuronal somata, based on their round morphological shape and their situation on top of an astrocytic monolayer. Smaller arrows indicate neurites. Scalebar in A applies for A-D. **(E-N)** Immunocytochemical labeling of the same culture at 61 days *in vitro*. Immunolabels include DNA-marker Hoechst (E, J; cyan), neuronal marker NeuN (F, K; magenta), reelin (G, L; orange) and oligomeric Aβ A11 (H, M; grey). Overlay of (E-H and J-M) is shown in (I and N), respectively; arrowheads indicate reelin-expressing neurons that co-expresses oligomeric Aβ. Scale bar in E applies for E-N. lLEC-LII; lateralmost lateral entorhinal cortex layer II, DIV; days in vitro.

### 3.2. Cultured lLEC-LII-neurons demonstrate spontaneous, sustained electrophysiological activity

An example of an MEA and its recording electrodes is shown in figure 4A. Both lLEC-LII neuron networks (MEA1 and MEA2) displayed spontaneous neuronal activity by 15 DIV, lasting until we terminated the cultures at 69 DIV (Fig. 4B). Their firing rate remained low throughout the whole recording period, and burst propensity was absent until about two months, when both networks showed a single burst (Fig. 4C).

**Figure 4.**
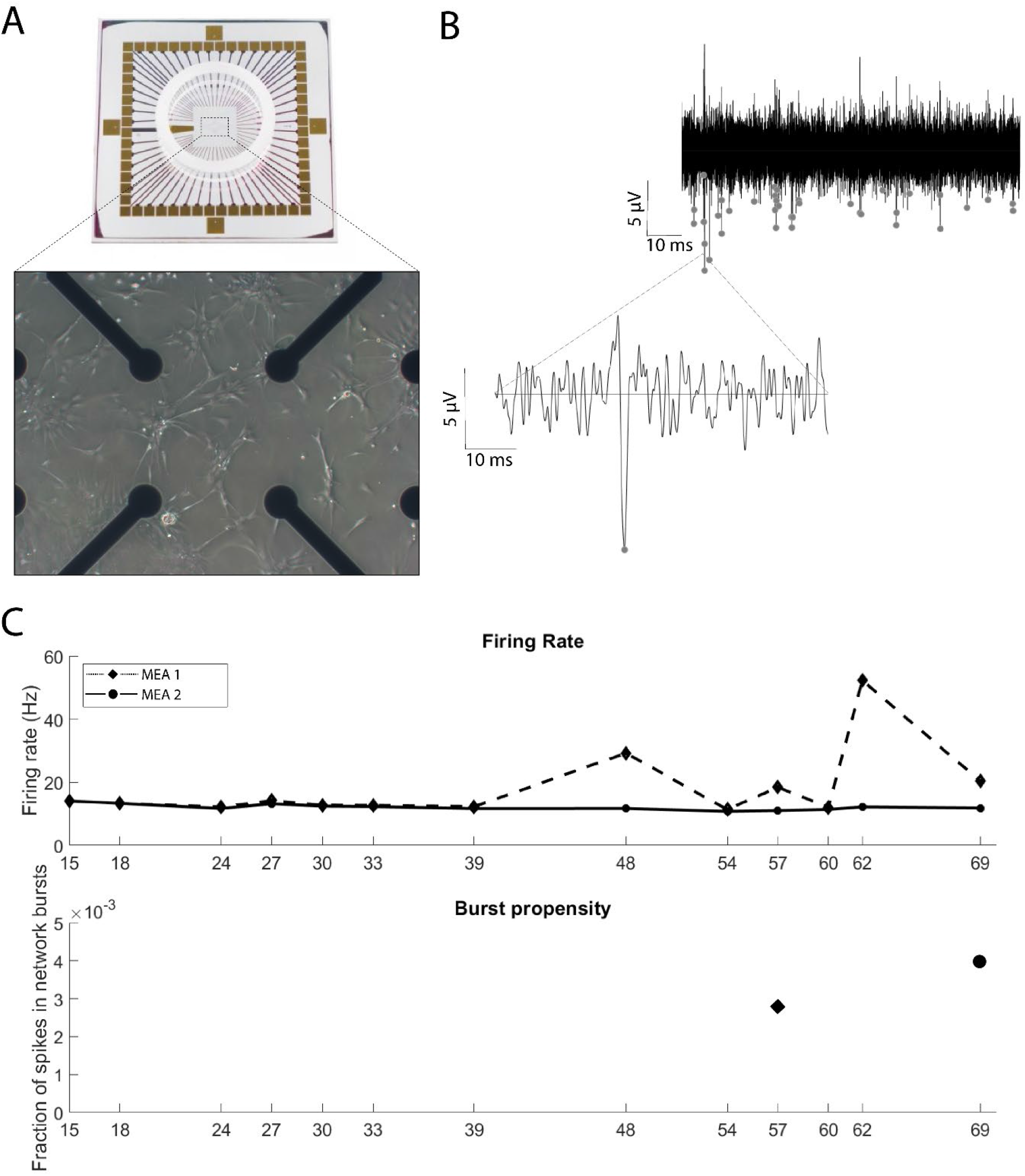
Cultured adult lLEC-LII neurons develop sustained electrophysiological activity. **(A)** lLEC-LII neuronal network at 24 DIV on an MEA. **(B)** Spike train (top) from one electrode with detected spikes indicated as grey dots. Inset (bottom) shows the waveform of an example spike. From recording at 33 DIV. **(C)** Firing rate (Hz) and burst propensity of the network from 15 DIV to 69 DIV. A single burst was detected for each culture as indicated (for MEA1 at 57 DIV; for MEA2 at 69 DIV). Abbreviations: lLEC-LII, lateralmost lateral entorhinal cortex layer II; DIV, days in vitro.

## 4. Discussion

To enable culturing of neurons and networks more relevant for the initiating stages of AD, we used adult APP/PS1 mice and microdissected tissue from the lateralmost domain of LEC, aiming for the Re+ lLEC-LII neurons. Once dissociated and plated, neurons self-organized into networks within two weeks and remained viable beyond two months (Fig. 3). We confirmed our ability to obtain the superficially situated Re+ lLEC-LII neurons by double-immunolabeling against NeuN and reelin, from which we observed several double-positive neurons (Fig. 3; Supplementary Fig. A.2). A further observation was that reelin expression co-localizes with that of Aβ, indicating that the tendency of such neurons to express Aβ (Kobro-Flatmoen et al., 2016) is retained when they are cultured (Fig. 3). In line with previous findings, an astrocytic feeder layer was crucial for survival and optimal growth of our cultured Re+ ECL-II neurons. More specifically, the presence of glial cells supports long term survival of adult primary neuronal cultures (Ray et al., 2009), promote synaptogenesis (Hama et al., 2004), and modulate neuronal excitability (reviewed in (Araque et al., 2004)).

By two weeks in culture, our neurons self-organized into networks and began exhibiting spontaneous spiking activity. The network complexity continued to increase over time, and the spontaneous spiking activity persisted for the entire culture period, consisting exclusively of very low burst propensity. This activity type may be a generic feature of adult cultured neurons, as previous studies reported that, in contrast to embryonic neuron cultures, adult neuron cultures tend to have non-repetitive firing patterns, rather exhibiting single-spike events (Evans et al., 1998; Varghese et al., 2009). Another possibility is that the low burst propensity we observe in our cultured lLEC-LII neurons is, at least in part, a result of the inherent biology of these neurons. Whole-cell current clamp experiments of lLEC-LII neurons revealed that they tend to fire late and with few spikes upon small depolarizing current-steps, and that a large depolarizing current-step is required to elicit a burst (Nilssen et al., 2018). The timing of media change relative to the time of recording is also likely to affect network activity. For example, human motor neurons were found to have a higher firing frequency 24 hours after media change compared to 48 hours after media change (Fiskum et al., 2021). The fact that we recorded activity 72 hours after media change may thus contribute to the very low burst propensity exhibited in our networks. An additional consideration is the fact that our lLEC-LII neurons express increased amounts of amyloid-β. Whether and how this affects lLEC-LII neurons was beyond the scope of this study and should be addressed in future work.

Aside from the relevance of Re+ lLEC-LII neurons in the onset of AD, it is likely that culturing of *adult* neurons offers a more relevant model for the disease, because it removes potential confounds owing to developmental factors operating in embryonic or early postnatal neurons. The ability to extract and move Re+ lLEC-LII neurons into an *in vitro* setting furthermore enables direct monitoring of their network properties and offers ease of access with respect to chemogenetic (Bauer et al., 2022; Westhaus et al., 2020), molecular-, (Valderhaug et al., 2021) and pharmacological tools (Han et al., 2017). Our approach may thus represent a valuable tool for studying very early proteinopathy and offers neuron- and network specific modelling of initiating AD pathology *in vitro*.

## Acknowledgements

We would like to thank PhD candidate Nicolai Winter Hjelm (INB, NTNU) and PhD candidate Vegard Fiskum (INB, NTNU) for help with Matlab scripts, and Hanne Mali Møllegård for breeding and genotyping of the APP/PS1 mice. This work was supported by a NTNU Enabling Technology Grant, the Olav Thon Foundation, Nasjonalforeningen for Folkehelsen, K.G. Jebsen Center for Alzheimer’s Disease, and Civitan Norwegian Research Fund for Alzheimer’s Disease.

## CRediT author statement

Katrine Sjaastad Hanssen; Methodology, Investigation, Formal analysis, Writing – original draft. Axel Sandvig; Resources, Writing – Review & Editing. Menno P. Witter; Resources, Writing – Review & Editing; Funding acquisition. Ioanna Sandvig; Resources, Conceptualization, Methodology, Data curation, Supervision, Funding acquisition, Writing – Review & Editing. Asgeir Kobro-Flatmoen; Conceptualization, Funding acquisition, Project administration, Supervision, Resources, Methodology, Data curation, Writing – Review & Editing.

## Appendix A

**Supplementary Table A.1.** Sex, age, and genotype for animals included in data analysis. DIV, Days in vitro; MEA, Microelectrode array; ICC, Immunocytochemistry.

| SEX | AGE | GENOTYPE | PURPOSE |
| --- | --- | --- | --- |
| M | P48 | APP/PS1 +/- | ICC 61 DIV |
| M | P48 | APP/PS1 +/- | ICC 61 DIV |
| M | P50 | APP/PS1 -/- | ICC 68 DIV |
| M | P50 | APP/PS1 -/- | ICC 68 DIV |
| M | P54 | APP/PS1 -/- | Nissl Staining |
| F | P60 | APP/PS1 +/- | Recordings (MEA1) |
| F | P60 | APP/PS1 +/- | Recordings (MEA1) |
| M | P60 | APP/PS1 +/- | Recordings (MEA2) |
| M | P60 | APP/PS1 +/- | Recordings (MEA2) |

DIV, Days in vitro; MEA, Microelectrode array; ICC, Immunocytochemistry.

**Supplementary figure A.1.**
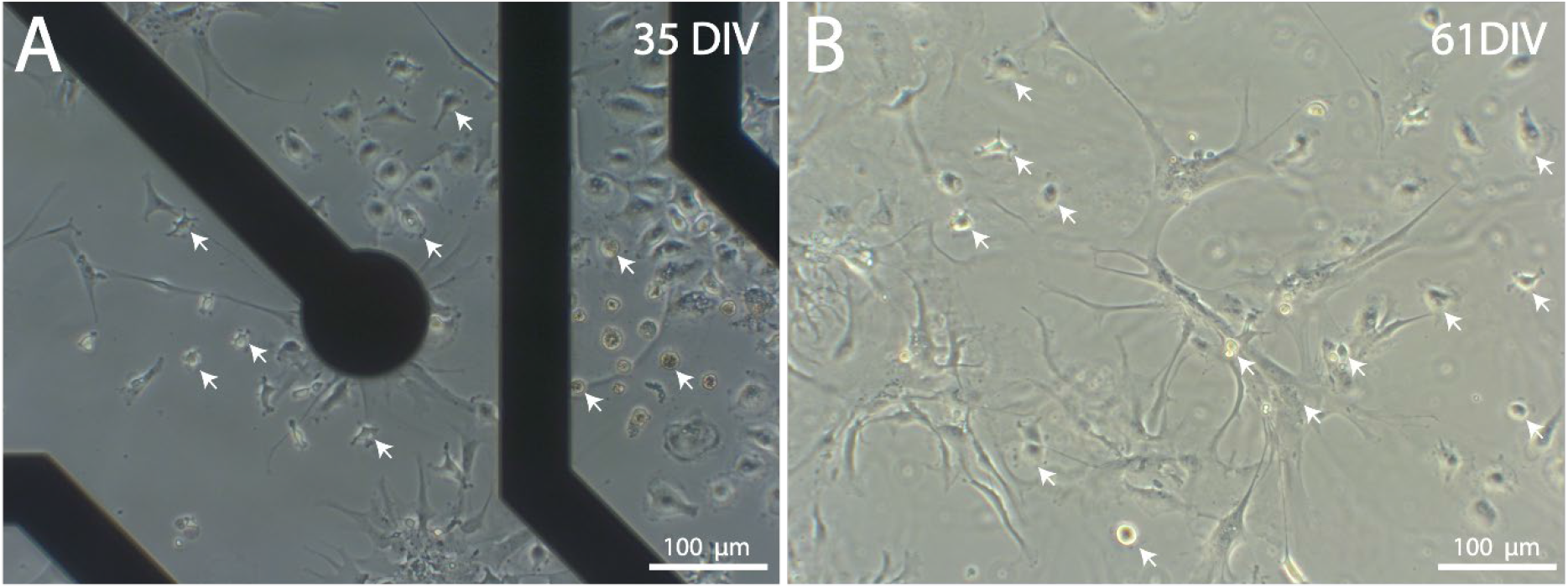
EC LII-neuron viability is dependent on frequency of media changes. **(A)** Lateralmost EC layer II neurons start to detach and die after 1 month with media changes conducted every 4 days. **(B)** EC layer II neurons start to detach and die after 2 months with media changes conducted every 3 days. Scale bars as indicated. DIV, Days in vitro.

**Supplementary Figure A.2.**
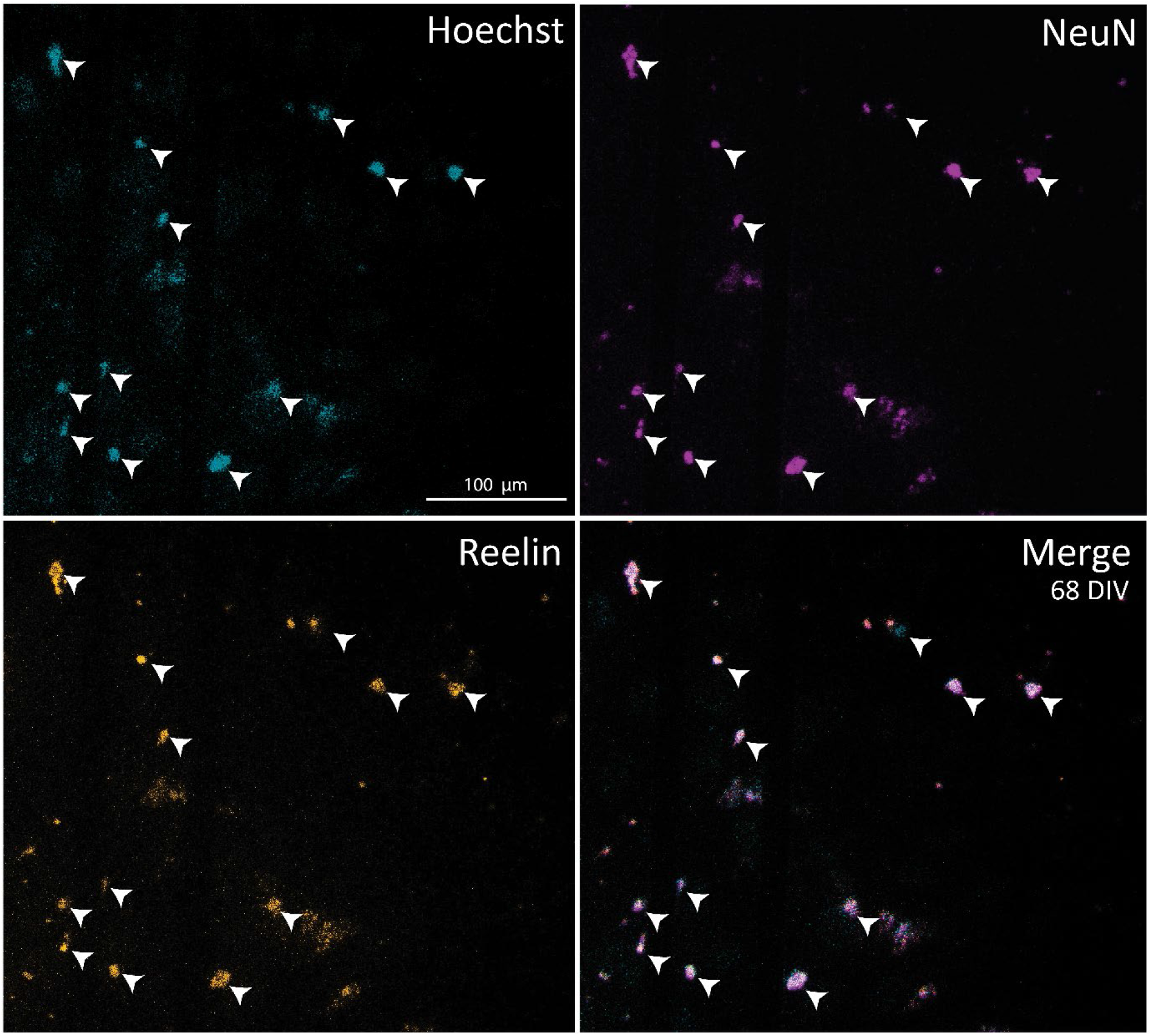
Culture of lLEC-LII neurons at 68DIV on a MEA. **A-D)** lLEC LII-neuron culture on a MEA at 68DIV. Staining of DNA-marker Hoechst (A; cyan), neuronal marker NeuN (B; magenta), and the glycoprotein reelin (C; orange) and Merge (D). Scale bars as indicated. lLEC-LII; lateralmost lateral entorhinal cortex layer II. DIV; days in vitro.

